# E-cigarette aerosol exposure of pulmonary surfactant impairs its surface tension reducing function

**DOI:** 10.1101/2022.08.16.501319

**Authors:** Emma Graham, Lynda McCaig, Gloria Shui-Kei Lau, Akash Tejura, Anne Cao, Yi Y. Zuo, Ruud Veldhuizen

**Author notes:** Emma Graham, (EG), Ruud Veldhuizen, (RV).

## Abstract

E-cigarette (EC) and vaping use continue to remain popular amongst teenage and young adult populations, despite several reports of vaping associated lung injury. This popularity is due in part to the vast variety of appealing flavours and nicotine concentrations easily accessible on the market. One of the first compounds that EC aerosols comes into contact within the lungs during a deep inhalation is pulmonary surfactant. This lipid protein mixture lines the alveoli, reducing surface tension and preventing alveolar collapse. Impairment of surfactant’s critical surface tension reducing activity can contribute to lung dysfunction. Currently, information on how EC aerosols impacts pulmonary surfactant remains limited. We hypothesized that exposure to EC aerosol impairs the surface tension reducing ability of surfactant. Bovine Lipid Extract Surfactant (BLES) was used as a model surfactant in a direct exposure syringe system. BLES (2ml) was placed in a syringe (30ml) attached to an EC. The generated aerosol was drawn into the syringe and then expelled, repeated 30 times. Biophysical analysis after exposure was completed using a constrained drop surfactometer (CDS). Minimum surface tensions increased after exposure to the EC aerosol. Variation in device used, addition of nicotine, or temperature of the aerosol had no additional effect. Two e-liquid flavours, menthol and red wedding, had further detrimental effects, resulting in higher surface tension than the vehicle exposed BLES. Alteration of surfactant properties through interaction with the produced aerosol was observed with a basic e-liquid vehicle, however additional compounds produced by added flavourings appeared to be able to increase inhibition. In conclusion, EC aerosols alter surfactant function through increases in minimum surface tension. This impairment may contribute to lung dysfunction and susceptibility to further injury.

## Introduction

Vaping and the use of e-cigarettes (ECs) has seen a marked increase over the last decade, particularly amongst youth and young adults (1–3). Although originally regarded as a popular smoking cessation tool, it is becoming increasingly clear that the target demographic for EC marketing has shifted from smokers to teenagers and young adults (3). Most of this population are not regular smokers, but instead are drawn to the attraction of efficient nicotine delivery and the large variety of appealing flavours available for recreational use (1, 3). Previous studies have indicated negative impacts of vaping (4). For example, in 2019 an outbreak of e-cigarette or vaping associated lung injury (EVALI), resulting in over 2000 hospitalizations and 68 deaths, was reported (5–7). Although studies associated the addition of vitamin E acetate with the incidences of EVALI, vaping without the compound has still been linked to lung injury, and there is not enough evidence to suggest vitamin E acetate to be the sole cause of injury (6, 8–10). Clearly, more information regarding the impact of vaping on the lung is required.

The process of vaping involves an EC device that uses a battery powered coil (atomizer) which heats an e-liquid comprised of solvents propylene glycol (PG) and vegetable glycerine (VG) to produce an inhalable aerosol. Many ECs utilize sub-ohm conditions, employing low resistance coils (blow 1 ohm) resulting in more rapid heating of the e-liquid and thicker more flavourful clouds. Complexity in studying the effects of this habit arise from the multitude of vaping options. There are presently over 16000 commercially available e-liquid flavour blends and almost 500 brands of ECs currently on the market (11–13). Adding to this complexity is the fact that many users make modifications to their device and/or generate their own e-liquid.

Furthermore, the harmful effects of vaping are not solely due to the impact of the ingredients of the e-liquid since it has been demonstrated that the generation of the aerosol results in the formation of a multitude of new products including formaldehyde and acetaldehyde (14, 15). This effect may vary among different devices, with particulate matter reported to increase with the evolution of some of the newer designs, thereby potentially increasing hazardous effects to the health of vapers (4, 16). Thus, although the general concept of vaping is relatively straightforward, complications in studying its effects arise from the variability and customization of the vaping process.

From a respiratory perspective, one of the first substances that EC aerosol encounters in the lungs is pulmonary surfactant. Surfactant is made up of 85-90% by weight phospholipids, 5-10% neutral lipids, mainly cholesterol, and 10% surfactant proteins, designated SP-A, SP-B, SP-C, and SP-D (17, 18). The primary function of surfactant is to reduce surface tension in the lung. To achieve this, surfactant rapidly absorbs to the surface forming the surface film, reaching a surface tension of ~23 mN/m, and subsequently reduces surface tension to ~0 mN/m upon compression (19, 20). This function is essential for breathing as it allows the lung to inflate and deflate with minimal changes in pressure, i.e., it maintains lung compliance, and prevents atelectasis (18, 21). As can be deduced from its function, absence or alterations to pulmonary surfactant can contribute to lung dysfunction. Previous studies have demonstrated that some ingredients of e-liquids can interfere with the function of model systems of surfactant (22–24).

Based on the above information, the objective was to investigate the impact that e-cigarette aerosol has on the function of surfactant. We initially focused on the impact of the common ingredients of aerosol, VG and PG, prior to testing the impact of selected additives. Using *in vitro* methods to replicate the dynamic compression and expansion of surfactant during respiration, this study focused on the biophysical changes to surfactant after direct exposure of a modified natural surfactant preparation to EC aerosol. It was hypothesized that EC aerosol inhibits surfactant’s ability to effectively reduce surface tension.

## Methods

### Materials

Bovine lipid extract surfactant (BLES) at 27mg/ml phospholipid concentration was obtained from BLES biochemicals inc (London, ON) and was diluted to 2mg/ml in a buffer containing 0.9% NaCl, 1.5mM CaCl2, and 2.5mM HEPES (pH 7.4). Three common vaping devices were used. The Aspire Pockex with 0.6 ohm coil, and Vaporesso Gen with GT4 0.15 ohm coil, were purchased from Vapour North (Pickering, ON). The Uwell Caliburn A2 with 0.9 ohm pre-installed mesh coil was purchased from Fogged Up Vape Shop (London, ON). All e-liquids were purchased from Vapour North.

### Exposure to aerosol

As schematically shown in figure 1, to expose surfactant to the EC aerosol, 2ml of BLES was placed in a 30ml syringe connected to an EC, via silicone tubing. The produced aerosol was drawn into the syringe and the syringe was placed on a rocker set to rotate 15-20 times/min for a 30 second exposure period. The aerosol was then expelled, and this procedure was repeated 30 times to simulate 30 inhalations. The control group consisted of BLES (2ml at 2mg/ml) placed in a 30ml syringe exposed only to air, no aerosol, and placed on the rocker for the same amount of time as the vaped samples. The exposed BLES was either used immediately for the analysis of surface tension reduction, or frozen at −20°C for subsequent analysis.

**Figure 1.**
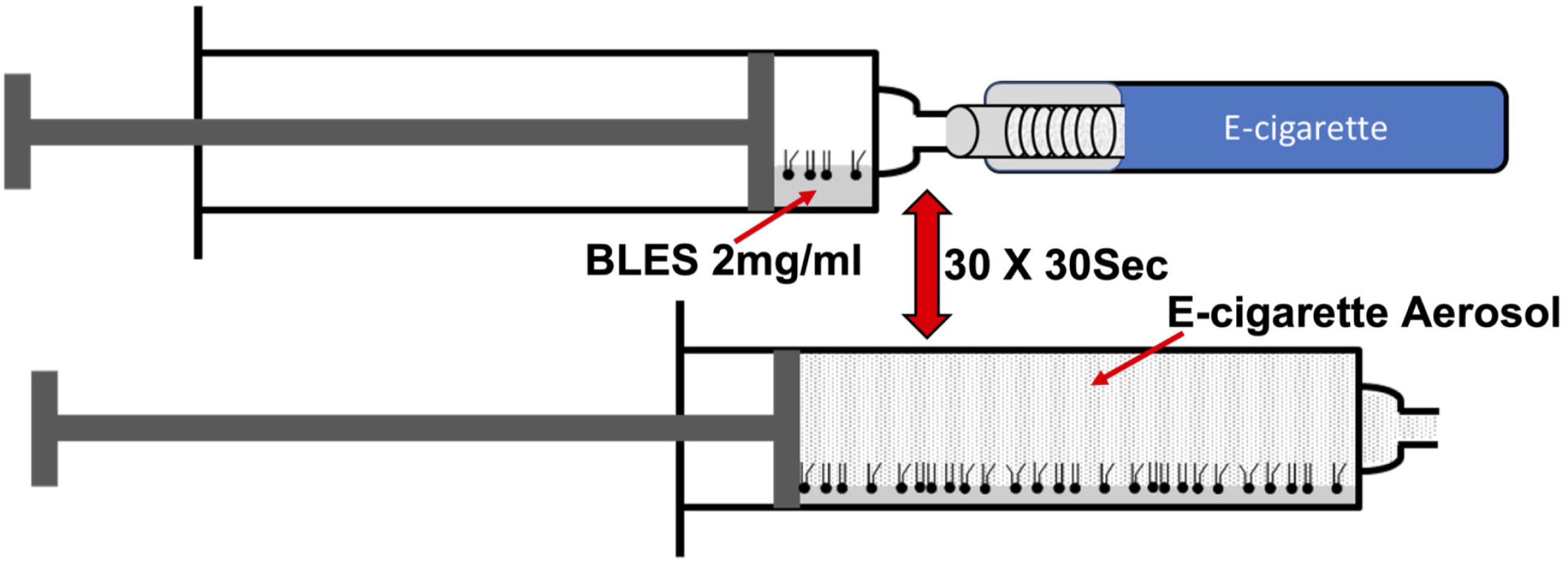
Schematic of the *in vitro* surfactant exposure set-up. Aerosol exposed surfactant was generated using a syringe containing BLES connected directly to an EC device which allowed aerosol to be drawn into the syringe and, after 30 seconds, expelled. This procedure was repeated 30 times.

### Biophysical analysis of surfactant

A constrained drop surfactometer (CDS) was used to assess the surface tension reducing properties of the BLES as previously described (25, 26). Briefly, an 8.5μL drop of the 2mg/ml surfactant sample was placed on the pedestal in an environmentally controlled chamber at 37 °C. A small capillary in the middle of the pedestal was attached to a computer controlled 2.5ml Hamilton syringe (Hamilton Co., Reno, NV) filled with distilled water. Two minutes after placing the BLES on the pedestal, the surface film of the drop was compressed and expanded via the controlled movement of the water in and out of the pedestal via the syringe. A total of 20 compression/expansion cycles were completed at a frequency of 1 cycle/ 1.5 seconds, with an area compression of approximately 20%. Aligned with the horizontal plane of the pedestal was a camera which took rapid pictures of the samples (15 fps) as they underwent dynamic changes. The images were then analysed by the Axisymmetric Drop Shape Analysis (ADSA) computer software to obtain precise measurements of the surface tension and surface area of the samples calculated based on the shape of the drop after the 2 minutes of adsorption as well as throughout the repeated compression/expansion cycles (27).

### Experimental design

Using the above procedure, a variety of exposure conditions were applied to surfactant followed by biophysical analysis. Initial experiments tested the e-liquid vehicle composed of unflavoured, nicotine free, e-liquid with a 50:50 VG:PG ratio using the Aspire Pockex. Subsequently, the 50:50 VG:PG liquid was tested in different devices, as well, other vehicle compositions (70:30 VG:PG and Max VG) tested with the Aspire Pockex device. The Vaporesso GEN EC was used to examine the effects of temperature through adjustment of the wattage in the device settings, with low (30W), medium (55W), and high (80W) temperatures. Of the three devices used, resistance varied between models (0.6ohm, 0.9ohm, 0.15ohm), and due to the simplicity of the Pockex and Caliburn models, atomizer temperature (through wattage) was only able to be assessed with the Vaporesso device. To determine the impact of aerosolization, vehicle 50:50 VG:PG e-liquid was mixed directly with BLES at 10 and 20% volumes, followed by analysis on the CDS.

To assess the impact of additives such as flavouring and nicotine to the vehicle, ten common e-liquids were tested: Vehicle + 18mg/ml nicotine, Juicy Burst + 3 mg/ml nicotine, RY4 Double (tobacco) + 18 mg/ml nicotine, RY4 Double (tobacco), Cigar Passion, Caramel Macchiato, Pink Lemonade, Red Wedding (raspberry doughnut with yogurt), Cinnaswirl, Menthol, Wintergreen, and Cherry. Experiments were performed in groups with multiple flavours, with each day of exposures including separate unexposed and vehicle exposed BLES controls.

### Statistics analysis

Data was expressed as the mean ± standard error of the mean (SEM) with a minimum of three independent replicates. Statistical analysis was performed using Graphpad Prism 9 software. Comparison of surface tensions across compression/expansion cycles for each sample were assessed using a two-way repeated measure analysis of variance (ANOVA) with a Bonferoni’s post hoc test. No significant differences between cycles were seen in samples except for the red wedding flavoured aerosol. Minimum surface tensions at the twentieth compression cycle were compared between groups using a one-way ANOVA, or two tailed unpaired t-test (air vs vehicle). For red wedding and menthol aerosols, minimum surface tensions at the first, fifth, tenth, and twentieth cycles were compared to vehicle and air control using a one-way ANOVA. Each set of three bars in Fig 4 comparing different flavourings, were analysed using a one-way ANOVA with a Dunnetts post hoc test compared to air control. P values less than or equal to 0.05 were considered statistically significant.

## Results

### Effect of vehicle aerosol on surfactant function

Using the above protocol, BLES was exposed to vehicle (50:50 VG:PG) aerosol and analyzed on the CDS. Representative surface tension versus relative area isotherms for the fifth and tenth compression/expansion cycle are shown in Figs 1A and 1B. At the fifth compression/expansion cycle, the air-exposed BLES sample shows a reduction in surface tension by compression to values below 5mN/m (lower curve) with subsequent expansion leading to an increase in surface tension with relative low amount of hysteresis (upper curve). In contrast, the vehicle exposed group had higher surface tension after compression (Fig 1A). Similar results were observed at the tenth cycle (Fig 1B). The results of the analysis of the minimum and maximum surface tension of all compression/expansion cycles are shown in Figs 1C and 1D. The air control BLES sample demonstrates low minimal surface tensions starting at the first compression that decrease further over the first few cycles to values below 5mN/m. Compared to these low values observed in the air control group, minimal surface tension in the vehicle exposed BLES group were significantly higher, reaching minimal values of approximately 10mN/m. Maximum surface tension of the control sample was approximately 37mN/m (Fig 1D), these values were not significantly different than those observed in the vehicle exposed group.

Additional data obtained from this analysis is shown in Table 1. The surface tension after two minutes of adsorption was not significantly different between the two groups. Furthermore, the calculated values for the maximum area compression, as well as the area compression required to reach minimum surface tension were also not significantly different between the two groups.

**Table 1.** Adsorption and compression data.

|  | Air Control | Vehicle Exposed |
| --- | --- | --- |
| Adsorption (mN/m) | 24.11 ±0.11 | 24.57±0.17 |
| Maximum Area Compression (%) | 20.33±0.21 | 20.65±0.73 |
| Area Compression at Min Surface Tension (%) | 19.27±0.31 | 19.41±0.90 |

Average adsorption and percent compression (±SEM) for air control and vehicle exposed groups

After establishing surfactant inhibition by the vehicle aerosol using a single experimental condition, this observation was further explored by examining two additional vehicle compositions, testing two other e-cigarette devices, examining three different wattages (i.e., temperatures) for aerosol generation, and mixing with non-aerosolized vehicle e-liquid. Fig 3A displays minimum surface tension trends (mN/m ± SEM) for BLES exposed to EC aerosol with each of the three different ratios of vehicle composition. Compared to air control samples, which reached low minimum surface tension values, 50:50 VG:PG vehicle achieved significantly higher minimum surface tensions. 70:30 VG/PG and max VG exposed samples were not significantly higher compared to air control. Minimum surface tensions obtained with aerosol from three different devices are shown in Fig 3B. Values for each of the devices were significantly higher than the air control. Comparison among the three EC devices revealed no statistically significant differences. A similar pattern was observed when comparing aerosol generated at three different wattages, as seen in Fig 3C. These wattages impact the temperature of the generated aerosol. The results show no difference among the varying wattages, with all three exposed samples resulting in significantly higher minimum surface tension than the air control. After mixing with e-liquid volumes up to 20%, no significant difference in minimum surface tension was observed compared to control, shown in Fig 3D. Surface tensions after two minutes of absorption, maximum area compression values, and maximum surface tensions were not significantly different between groups and are shown in supplementary data (S1 and S2).

**Fig 2.**
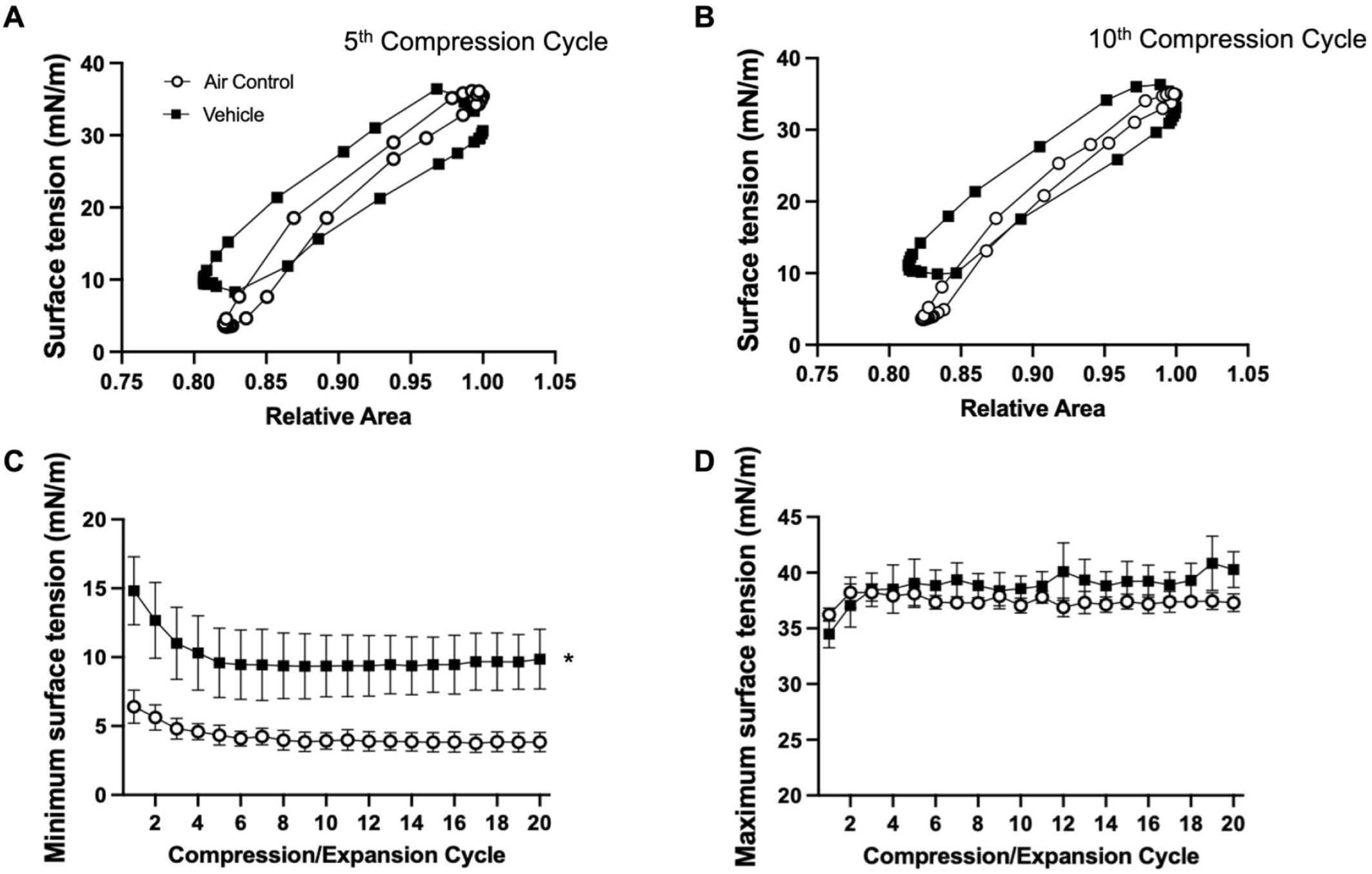
Vehicle compression/expansion. BLES exposed to vehicle (■) compared to air control (○). **A.** Representative surface tension vs area isotherm at the fifth compression/expansion cycle. **B.** Representative surface tension vs area isotherm at the tenth compression/expansion cycle. **C.** Minimum surface tensions across 20 compression/expansion cycles. **D.** Maximum surface tensions across 20 compression/expansion cycles. *P<0.05 compared to air control.

**Fig 3.**
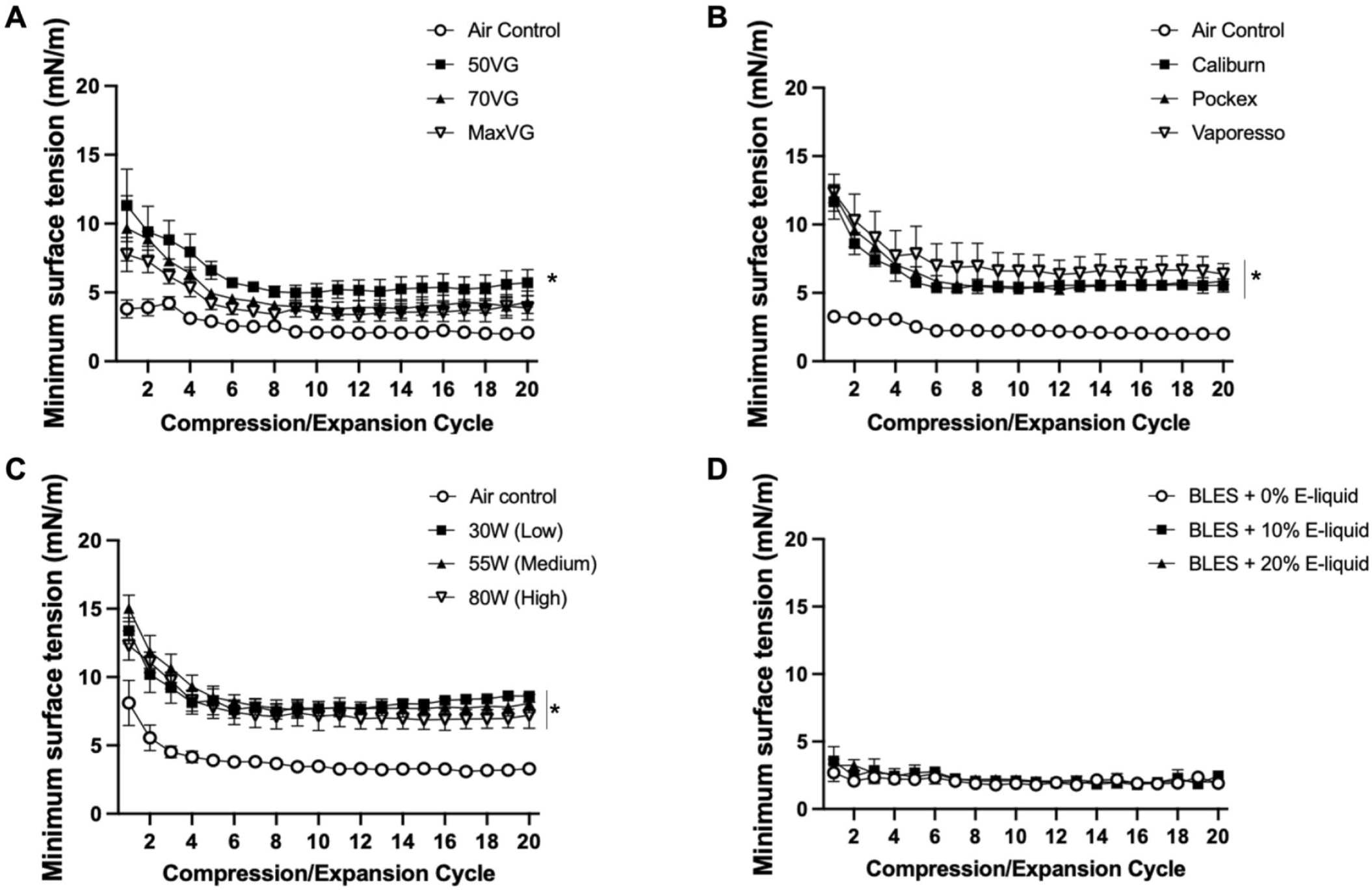
Minimum surface tensions of experimental variables. Minimum surface tensions (± SEM) across 20 compression/expansion cycles for BLES exposed to three experimental variables, compared to air control. **A.** Three different vehicle compositions with increasing VG content. **B**. Three different EC devices. **C.** Three different wattages set on a programmable EC to alter aerosol temperature. **D.** BLES mixed with three different volumes of vehicle e-liquid. N=3 *P<0.05 compared to air control.

### Effect of commercially available e-liquid aerosols

Having established the effects of the vehicle on surfactant, our subsequent experiment evaluated the effect of aerosols from a variety of e-liquids containing flavourings and/or nicotine. Ten popular commercially available e-liquids were tested. Fig 4 summarizes the data by displaying the minimum surface tension values obtained for the fifth compression cycle as compared to its vehicle and air control. The values for each of the 20 compression/expansion cycles are shown in the supplementary data (S2 Fig. A-I). Consistent with the above experiments, in each of the tests the vehicle exposure led to higher minimum surface tension as compared to air-exposed BLES (Fig 4). With the exception of two of the e-liquids, all flavours resulted in minimum surface tension that were not significantly different to the unflavoured vehicle. This included samples with nicotine additives. The two exceptions to this pattern were menthol and red wedding flavoured aerosols, which resulted in significantly increased minimum surface tension compared to both air control and the unflavoured vehicle control.

**Fig 4.**
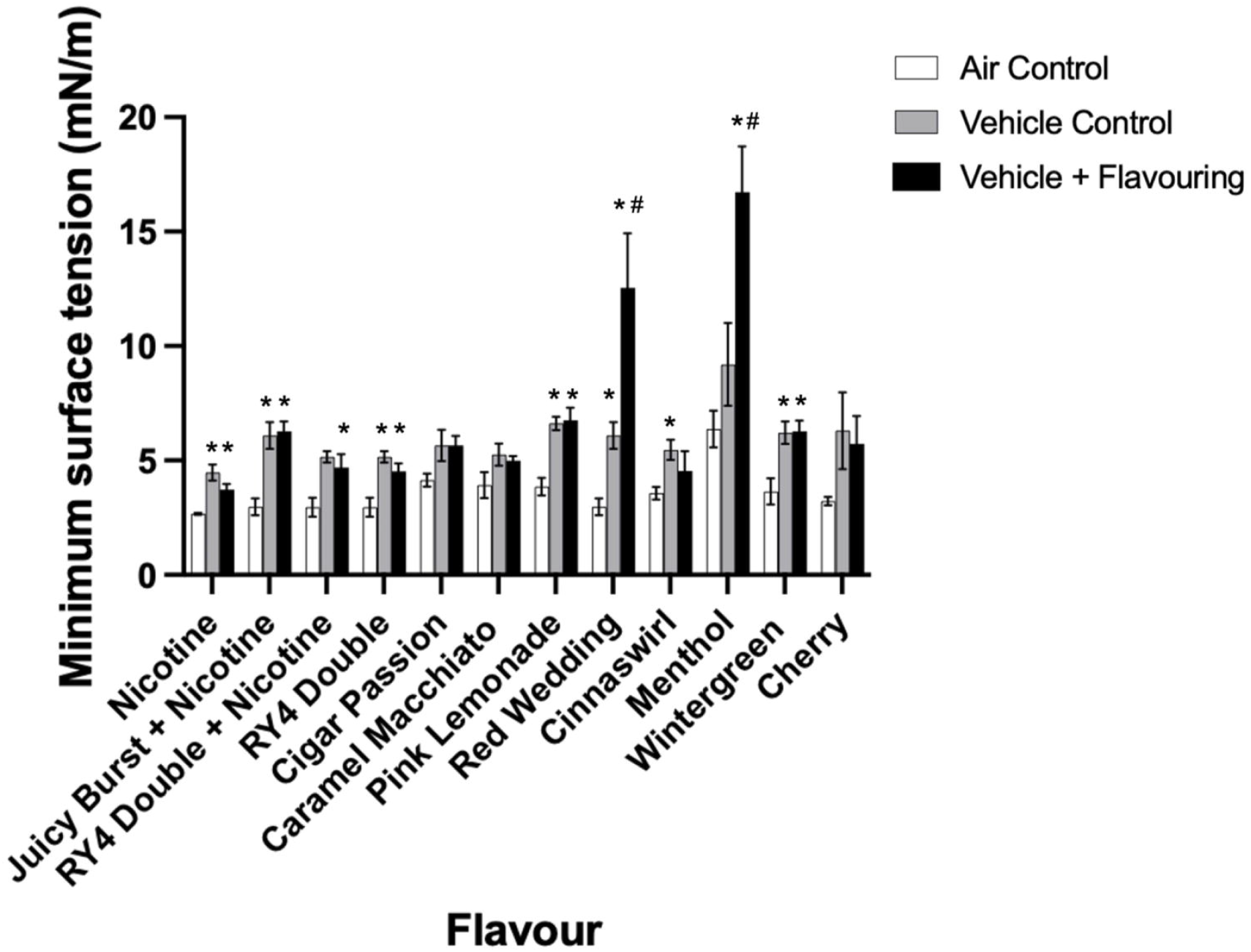
Minimum surface tensions of flavoured e-liquids. BLES exposed to aerosols containing nicotine and/or flavouring additives compared to air control and unflavoured vehicle at the fifth compression/expansion cycle (± SEM). Each flavour on the x-axis represents BLES exposed to air, unflavoured vehicle, and vehicle + flavouring aerosols. N=3-4 *P<0.05 compared to air control, #P<0.05 compared to vehicle.

A further exploration of the data obtained from menthol and red wedding e-liquids is shown in Fig 5 (Menthol) and Fig 6 (Red Wedding). Isotherms for the fifth (Fig 5A) and tenth (Fig 5B) cycles show menthol exposed BLES to result in minimum surface tension at ~20 mN/m, higher than the vehicle exposed BLES which achieved minimum surface tension of ~10 mN/m. Menthol exposure resulted in significantly higher minimum surface tension than both unflavoured vehicle and air control groups at all 20 compression/expansion cycles (Fig 5C). Maximum surface tension remained consistent between groups, at ~35 mN/m (Fig 5D). The inhibitory effect of menthol on minimum surface tension was further enhanced with three repeated exposures on consecutive days (S4). Isotherms at the fifth cycle for red wedding exposed BLES, also show significant impairment, with minimum surface tension at ~20 mN/m, higher than both the vehicle and air exposed groups (Fig 6A). At the tenth cycle however, the red wedding flavour exposed group achieves minimum surface tension similar to that of the vehicle exposed group (Fig 6B). A steep decline on a cycle-by-cycle basis can be observed for the red wedding exposed group, until approx. the tenth cycle, where surface tension remains similar to the vehicle exposed (Fig 6C). Both flavoured and vehicle exposed BLES showed higher minimum surface tension than the air control. Red wedding further appeared to have lower maximum surface tension during the first ten cycles, maintaining values ~30 mN/m, whereas air control groups had an average maximum surface tension of ~40 mN/m (Fig 6D).

**Fig 5.**
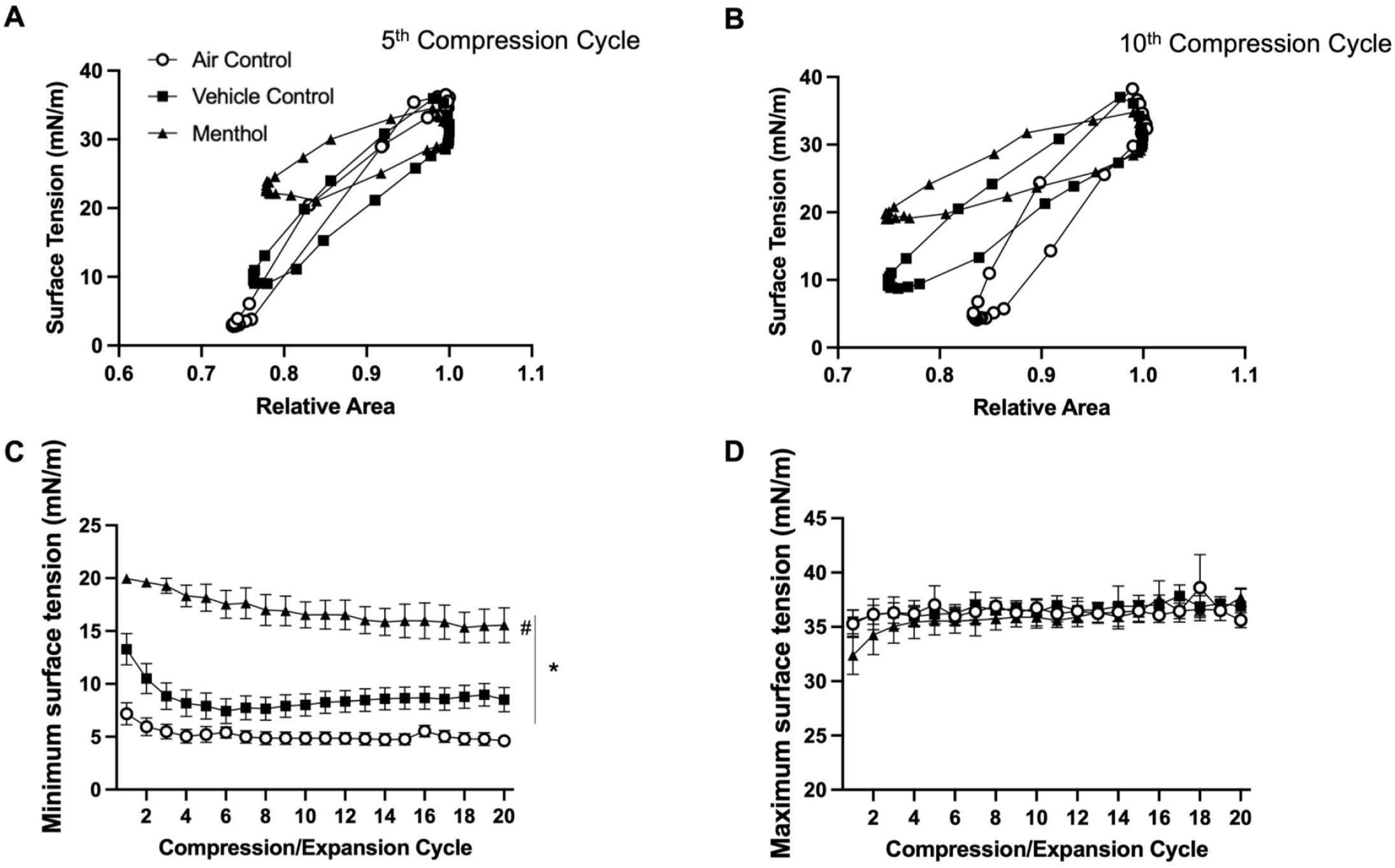
Menthol compression/expansion data. BLES exposed to menthol flavoured aerosol (▲) compared to vehicle (■) and air control (○). **A.** Representative surface tension vs area isotherm at the fifth compression/expansion cycle. **B.** Representative surface tension vs area isotherm at the tenth compression/expansion cycle. **C.** Minimum surface tensions (± SEM) across 20 compression/expansion cycles. **D.** Maximum surface tensions (± SEM) across 20 compression/expansion cycles. (N=5) *P<0.05 compared to air control, #P<0.05 compared to vehicle.

**Fig 6.**
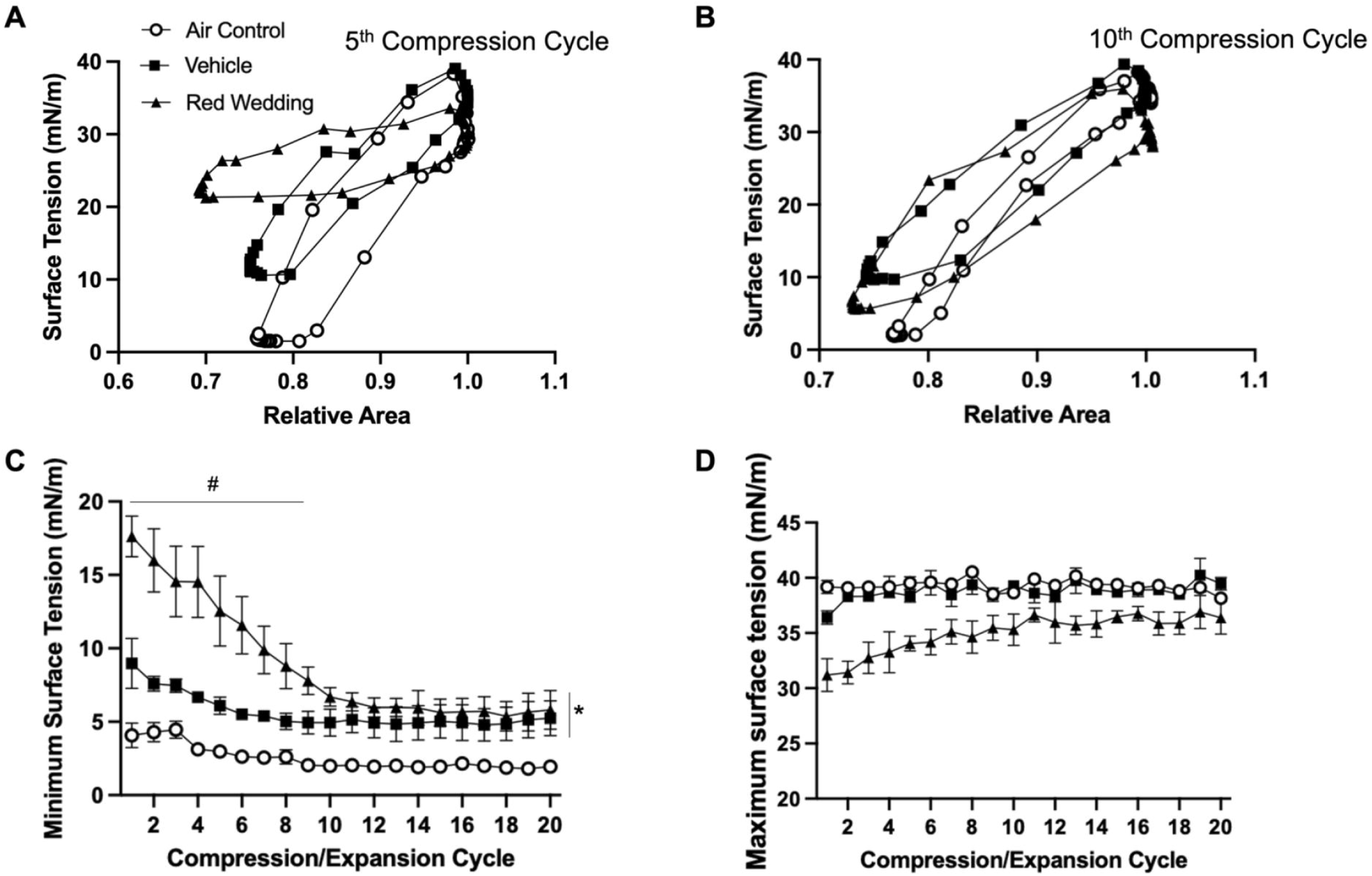
Red wedding compression/expansion data. BLES exposed to red wedding flavoured aerosol (▲) compared to vehicle (■) and air control (○). **A.** Representative surface tension vs area isotherm at the fifth compression/expansion cycle. **B.** Representative surface tension vs area isotherm at the tenth compression/expansion cycle. **C.** Minimum surface tensions (± SEM) across 20 compression/expansion cycles. **D.** Maximum surface tensions (± SEM) across 20 compression/expansion cycles. (N=3) *P<0.05 compared to air control, #P<0.05 compared to vehicle.

## Discussion

This study tested the hypothesis that EC aerosols inhibit surfactant’s ability to effectively reduce surface tension. Results supported this hypothesis as exposure to EC aerosols resulted in impaired surfactant function, in particular its ability to reduce surface tension to near-zero values. All exposed samples, regardless of additive, device, or vehicle composition resulted in higher minimum surface tensions than air exposed samples. Furthermore, surfactant dysfunction was observed in all cases of aerosol exposure with different additives, and most of the flavours tested resulted in similar inhibition and minimum surface tensions to those of the vehicle. Interestingly, two flavours, menthol and red wedding, resulted in significantly higher minimum surface tensions compared to the vehicle. It is concluded that aerosol exposure can lead to surfactant impairment which, *in vivo*, could impact lung function. This effect of EC aerosol can be augmented due to certain flavour additives.

Previous studies have examined the effects of e-liquid aerosols on surfactant, with a variety of experimental approaches. Distinctions in the type of surfactants used, with several studies using surfactant-like lipid mixtures, as well as differences in exposure method including mixing of surfactant with isolated components of the e-liquid directly, have been reported (22, 28, 29). Furthermore, a variety of analyses, including structural and functional assessments have been reported (24, 30). Building on these studies we used a modified natural surfactant, exposed the samples to EC generated aerosols, and focused on the main function of surfactant by measuring surface tension reduction. Specifically, we used BLES, a well-established commonly prescribed exogenous surfactant composed of the surfactant lipids and the hydrophobic proteins B and C (31). Aerosol exposure was performed in a syringe containing a surfactant suspension allowing direct exposure of surfactant to the aerosol. This exposure of the surfactant before compression/expansion cycling resulted in findings consistent with recent direct aerosol exposure of a compressing drop (30). Analysis was done using the CDS, which was able to provide extremely accurate measurements of changes in surface tension and droplet area during consistent dynamic cycling with compression maintained within physiological levels (19). To provide a broad assessment of the impact of EC aerosols on surfactant, we used several commercially available EC devices and multiple e-liquids to accurately reflect the vaping tools on the market today. Overall, we deemed that this approach allowed for effective testing of our hypothesis.

One of the main goals of our study was to examine the effect of the base vehicle of e-liquid, VG and PG, since these compounds are universally utilized during vaping. Although both compounds are safe and frequently used humectants in a wide range of foods and consumer cosmetic products (32–34), research on the effects of aerosolization and inhalation of these compounds is limited. Focusing on the effect of surfactant specifically, it has been previously shown that EC aerosols did not induce harmful effects on surfactant interfacial properties but caused minor alterations in lateral structure arrangement after exposure (24). In this case however, surfactant (Infasurf) was not directly exposed to the aerosol, but rather the aerosol was bubbled into the subphase in a Langmuir-Blodgett trough before addition of the surfactant, without measurement of aerosol concentrations. Alterations observed to the lateral structure are nonetheless important to note, as proper arrangement of the surface film phospholipids are essential to reaching near zero surface tensions upon compression (35). In two other studies, impairment of surfactant function was observed. Using a lipid mixture as a model system of surfactant it was observed that exposure to EC aerosol resulted in lower maximum surface pressures at any given area (higher minimum surface tensions) and altered compressibility on a Langmuir balance (28). Further, direct mixing of non-aerosolized PG and VG with surfactant, resulted in impaired surfactant function (increased surface tension amplitude, mean surface tensions, and surface elasticity), particularly in ultra-high concentrations of PG dominant mixtures (29). The current study builds upon these findings, by exposing a natural surfactant to aerosolized VG/PG and dynamic measurement of surfactant function. Furthermore, we showed that the VG/PG induced reduction in surface activity was consistent among several different sub-ohm devices with very low resistance coils and different wattages for aerosol generation. Although the sub-ohm devices are popular amongst users, large variation in resistance and the amount of current passed through the coil exist beyond those tested in the current study. Overall, the findings support a modification in dynamic surface-active properties of surfactant due to vehicle aerosol exposure.

Having established a general, albeit mild, inhibition of surfactant by EC aerosol in general, our second goal was to examine the effect of different flavored e-liquids. Rationally selecting specific e-liquids among the approximately 16000 flavours currently available is difficult, however, we picked popular e-liquids in different categories (tobacco alternatives, drinks, candy, and dessert) (13). Interestingly, whereas most e-liquids showed similar results to vehicle, menthol and red wedding resulted in further inhibition of surfactant function. The mechanism of inhibition of surfactant by menthol has been recently studied and indicates a direct interaction of this flavouring with surfactant phospholipids and proteins thereby impacting the surface tension reducing surface film (30). The specific process by which red wedding flavouring impacts surfactant is not clear, and studies of the lateral organization of the surfactant film, for example by atomic force microscopy could explore this further. Nevertheless, the pattern of inhibition in which high surface tensions are only observed in the early compression/expansion cycles would suggest that chemicals are inserted into the surface film but are removed, or squeezed out, during repeated compression and expansion (19, 36). In contrast, it appears as if surfactant remains inhibited by menthol exposure across compression/expansion cycles, suggesting a more permanent alteration of the surface film. A recent study, showing similar effects to menthol as the current study, did indeed show such alteration of the surface film using atomic force microscopy (30). Regardless of the specific mechanism, the data clearly indicate that variability exist between the impact of aerosols from different e-liquids. Unlike in food and beverages, the chemicals in e-liquids are not highly regulated, and not all the specific ingredients are disclosed. For example, a study examining the chemical concentrations in JUUL pod-based e-liquids identified 59 chemicals, with menthol, vanillin, and ethyl maltol having the highest concentrations, with the remaining 56 chemicals having concentrations of <1.0mg/mL (37). Considering our data demonstrated enhanced surfactant inhibition with 2 out of 12 e-liquid samples selected, we speculate that a substantial fraction of the commercially available e-liquid flavours available have a negative impact on surfactant function.

Further complicating the study of EC aerosol interactions with the surfactant system is the generation of various compounds during the aerosolization process. For example, flavour aldehydes have been shown to react causing the formation of PG acetals (38). Benzaldehyde (cherry flavouring), ethylvanillin, vanillin, cinnamaldehyde, and citral were examined to determine reaction products after vaping and were shown to produce PG acetals that remained stable for hours (38). It has also been reported that aerosolization, and subsequent degradation of VG/PG can lead to the generation of harmful chemicals, including carbonyl compounds such as formaldehyde, acetaldehyde, and acrolein, as well as volatile organic compounds, tobacco-specific nitrosamines, metals, free radicals, and reactive oxygen species in EC aerosols (15, 39–41). In a non-targeted study, over 115 volatile organic compounds were found in a single puff of EC aerosol (42). Further, compared to the composition of the non-aerosolized liquid, aerosolization produced more than 80 additional compounds beyond the 60 identified within the liquid itself (42). It has also been reported that higher power EC devices increase reactive oxygen species production (43). However, it should be noted that in the current study, no differences were observed between three devices, as well as no significant differences between power levels (wattage) on our highest power device (Vaporesso), suggesting that either the lowest wattage was able to maximally effect the surface tension of the surfactant, or that reactive oxygen species production was not a major contributor to the observed surfactant inhibition. Furthermore, no significant differences were seen when mixing the non-aerosolized e-liquid directly with BLES, indicating the importance of aerosolization for inhibition. Overall, the published data suggest that the inhibition of surfactant by EC aerosols may not only be the results of the incorporation of e-liquid components but also due to additional components generated during aerosolization. This includes those generated from flavour additives, which can either insert into the surfactant and/or chemically react with surfactant lipids and proteins. As such, it is important that future regulations regarding vaping and e-liquid composition should base recommendations on studies that have evaluated the health impact after aerosol generation.

Although the current study examined the effects of EC aerosols on surfactant *in vitro*, the rationale for our experiments was the interaction of aerosols with surfactant in the lung, thus it is important to extrapolate our findings to the *in vivo* situation. Although we observed a slight decrease in surface tension reducing ability after exposure, such minor surfactant dysfunction may not have a major impact *in vivo*. Nevertheless, a clinical study by Dr. Vardavas and colleagues demonstrated that using an EC device for only five minutes led to an immediate, subtle, decrease in lung mechanics in a group of volunteers (44). Considering the rapid nature of this effect on lung mechanics, it would be consistent with a rapid direct interaction of the aerosol with surfactant, rather than, for example, a cellular mechanism. A second potential consequence of altered surfactant function may be a susceptibility to lung injury. Thus, although vaping induced alterations to surfactant may not be sufficient to cause lung dysfunction, in scenarios where the surfactant is further challenged it may become more impaired and ultimately affect lung function. For example, it has been shown that surfactant that contains lyso-phospholipids was more susceptible to inhibition by serum proteins as compared to control (45). It should also be noted that *in vivo* vaping may impact other aspect of surfactant, such as the expression of the surfactant associated proteins. Two previous studies show different results in this regard with one study observing dysregulation of certain genes, but no significant alterations to surfactant proteins A, B, C, or D (46), but another reporting decreased expression of surfactant proteins (23). Difference in experimental set up, specific vaping conditions and exposure times may have contributed to these differences. Future studies examining the acute impact of EC aerosols on surfactant *in vivo*, as well as susceptibility of aerosol exposed surfactant to further inhibition, are warranted.

Our study has several limitations. First the current study uses an exogenous surfactant, BLES, to test the impact of EC aerosol. All studies on the effect of vaping have utilized either exogenous surfactant or an even more limited, lipid model system. The rationale for using exogenous surfactant is that it contains all surfactant lipids and the hydrophobic proteins, SP-B and C, and has consistent functional properties. Nevertheless, future studies with natural surfactants from animal and/or humans would provide additional information of the impact of the hydrophilic surfactant protein A, as well as other components that co-isolate with surfactant obtained from lung lavage. A second limitation is that nicotine was only tested individually as well as with two flavour combinations. As no additional affect was observed, we neglected to test nicotine further, however it is possible that nicotine could have an interaction with specific flavours to potentially affect the impact on surfactant. Lastly, only the effects of acute exposure on surfactant were tested. People that vape will do so in a prolonged fashion which may have additional consequences. For example, previous *in vivo* work on mice exposed daily to EC aerosols for several weeks showed altered lipid homeostasis in alveolar macrophages, as well as accumulation of phospholipids in bronchoalveolar lavage fluid (23). The impact of EC aerosols on surfactant *in vivo* may thus be affected by acute direct effect as observed in the current experiments, as well as prolonged effects related to the impact on surfactant synthesis and metabolism.

## Conclusion

This study demonstrated that the aerosol produced from the e-liquid vehicle (VG/PG) was able to alter surfactant function independent of nicotine or device style, with some flavourings providing further inhibition. From these results we conclude that acute EC aerosol exposure leads to impairment of the pulmonary surfactant system through increases in minimum surface tension. We speculate that this has potential detrimental effects in the *in vivo* scenario by contributing to lung dysfunction and making the lung more susceptible to further injury.

## Acknowledgments

We would like to thank Dr Fred Possmayer and Dr Cory Yamashita for helpful discussion, as well as Justin Rao, Sabrine Gehani, and Alexandra Troitskaya for support and assistance in running experiments. Help with the statistical from Dr Eric Patterson is greatly appreciated. The exogenous surfactant, BLES, was a generous gift from BLES biochemicals (London On. Canada).

## Supporting information

**S1 Fig. A-D. Maximum surface tensions of experimental variables.** Maximum surface tensions (± SEM) across 20 compression/expansion cycles for BLES exposed to three experimental variables, compared to air control. **A.** Three different vehicle compositions with increasing VG content. **B**. Three different EC devices. **C.** Three different wattages set on a programmable EC to alter aerosol temperature. **D.** BLES mixed with three different volumes of vehicle e-liquid. N=3 *P<0.05 compared to air control.

**S2. Table.**
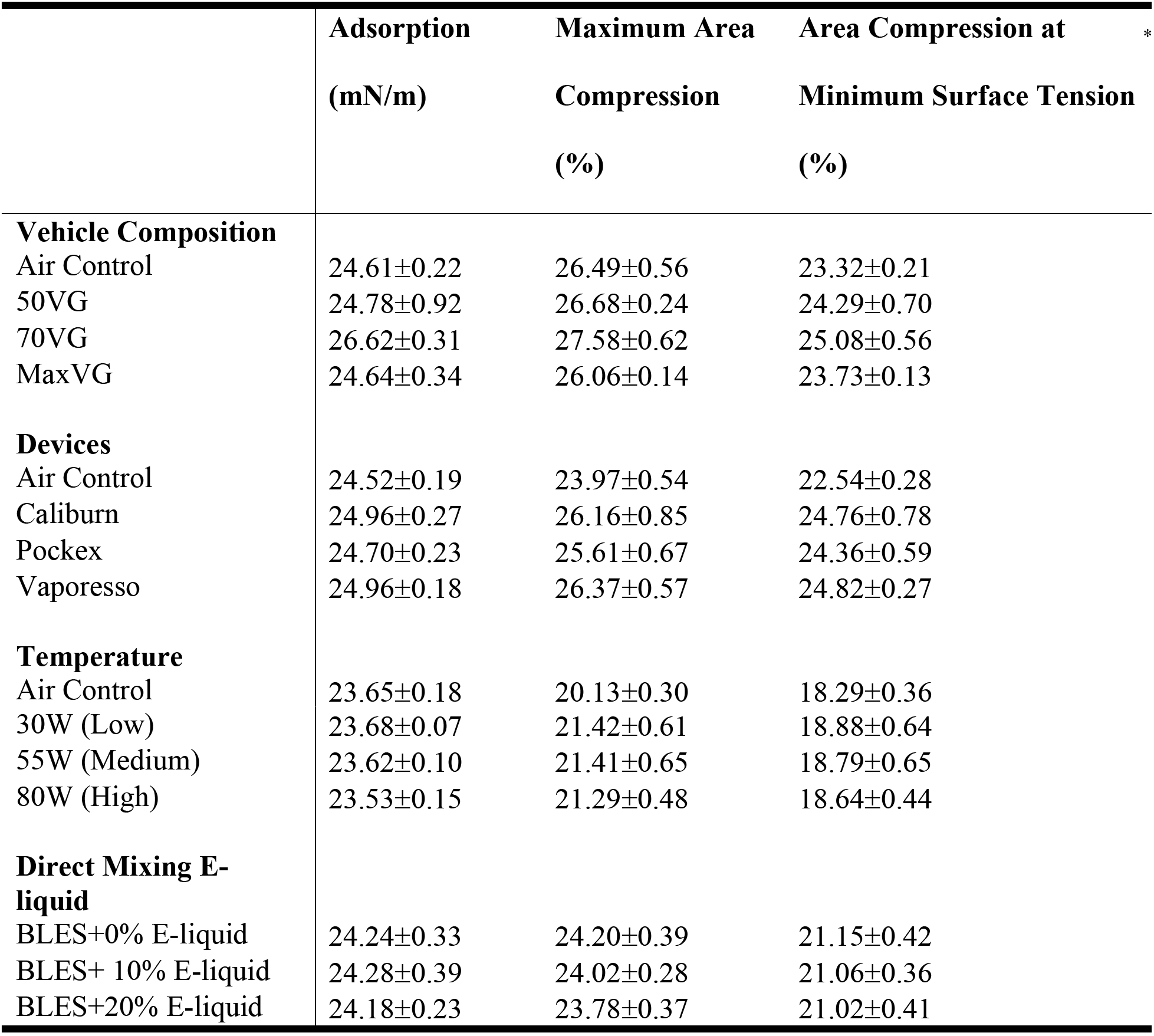
Adsorption and compression data. Average adsorption and percent compression (±SEM) for control and exposed groups under four experimental conditions: changing vehicle composition, using different EC devices, varying temperature through increasing wattage, and mixing of non-aerosolized e-liquid directly with BLES.

|  | Adsorption<br>(mN/m) | Maximum Area<br>Compression<br>(%) | Area Compression at<br>Minimum Surface Tension<br>(%) | * |
| --- | --- | --- | --- | --- |
| **Vehicle Composition** |  |  |  |  |
| Air Control | 24.61±0.22 | 26.49±0.56 | 23.32±0.21 | |
| 50VG | 24.78±0.92 | 26.68±0.24 | 24.29±0.70 | |
| 70VG | 26.62±0.31 | 27.58±0.62 | 25.08±0.56 | |
| MaxVG | 24.64±0.34 | 26.06±0.14 | 23.73±0.13 | |
| **Devices** |  |  |  |  |
| Air Control | 24.52±0.19 | 23.97±0.54 | 22.54±0.28 | |
| Caliburn | 24.96±0.27 | 26.16±0.85 | 24.76±0.78 | |
| Pockex | 24.70±0.23 | 25.61±0.67 | 24.36±0.59 | |
| Vaporesso | 24.96±0.18 | 26.37±0.57 | 24.82±0.27 | |
| **Temperature** |  |  |  |  |
| Air Control | 23.65±0.18 | 20.13±0.30 | 18.29±0.36 | |

|  |  |  |  |
| --- | --- | --- | --- |
| 30W (Low) | 23.68±0.07 | 21.42±0.61 | 18.88±0.64 |
| 55W (Medium) | 23.62±0.10 | 21.41±0.65 | 18.79±0.65 |
| 80W (High) | 23.53±0.15 | 21.29±0.48 | 18.64±0.44 |
| **Direct Mixing E-liquid** |  |  |  |
| BLES+0% E-liquid | 24.24±0.33 | 24.20±0.39 | 21.15±0.42 |
| BLES+ 10% E-liquid | 24.28±0.39 | 24.02±0.28 | 21.06±0.36 |
| BLES+20% E-liquid | 24.18±0.23 | 23.78±0.37 | 21.02±0.41 |

**S3 Fig. A-I.** Minimum surface tensions (± SEM) across 20 compression/expansion cycles for BLES exposed to nine flavourings/additives compared to unflavoured vehicle control. N=3.

**S4 Fig.** Minimum surface tensions (±SEM) across 20 compression/expansion cycles for BLES exposed to menthol and vehicle aerosols as well as air control after one day and three days of consecutive exposure. N=3 *P<0.05 compared to day one.

