## Supplementary figures and images for "E-cigarette aerosol exposure of pulmonary surfactant impairs its surface tension reducing function"

### Supplemental Figure 1

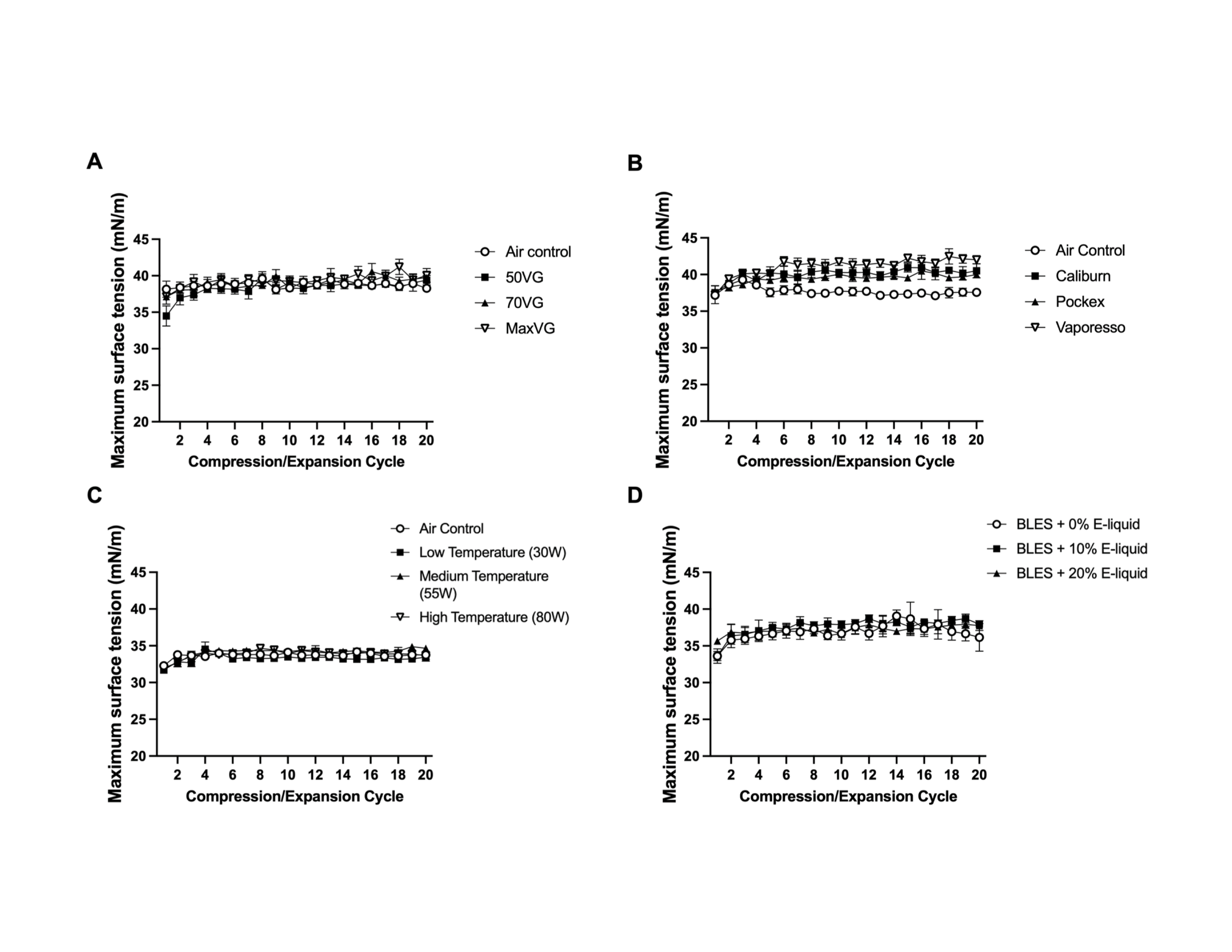

### Supplemental Figure 2

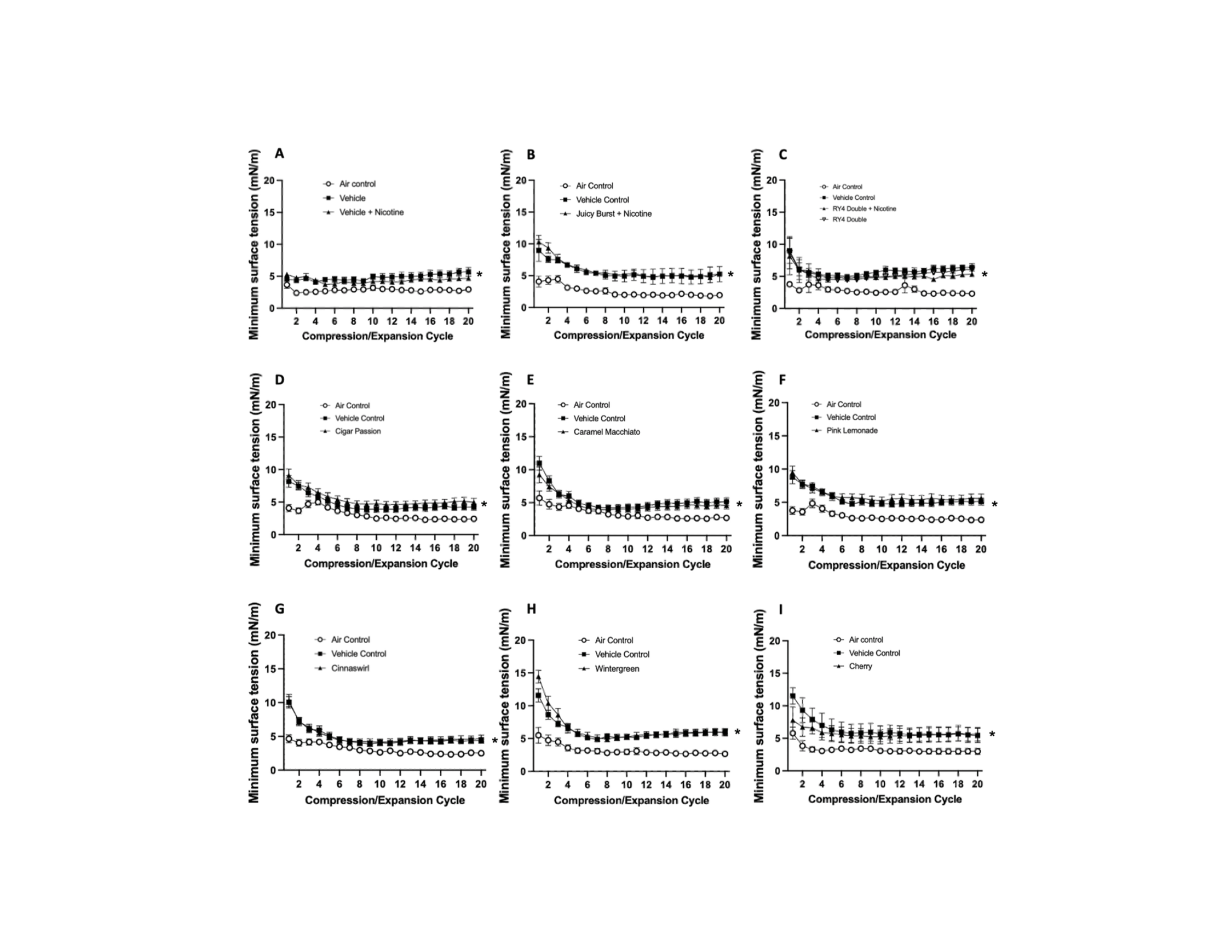

### Supplemental Figure 3

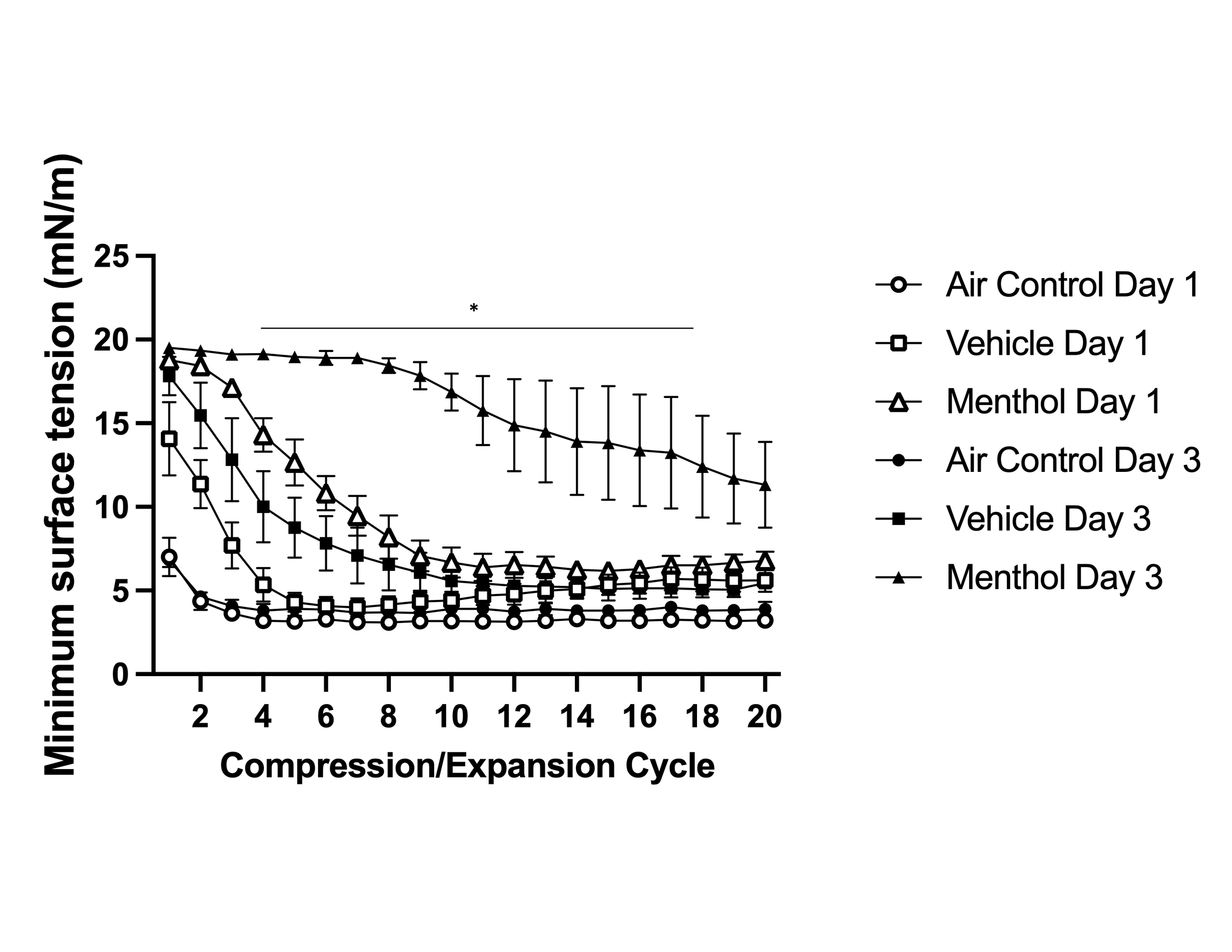
